# Force-dependent stabilization of apical actomyosin by Lmo7 during vertebrate neurulation

**DOI:** 10.64898/2026.01.24.701484

**Authors:** Miho Matsuda, Sergei Y. Sokol

**Affiliations:** Department of Cell, Developmental and Regenerative Biology, Icahn School of Medicine at Mount Sinai, New York, NY, USA

## Abstract

Coordinated control of actomyosin contractility is essential for epithelial morphogenesis, including vertebrate neural tube closure. Lim domain only 7 (Lmo7) is a force-sensitive regulator of contractility that binds non-muscle myosin II (NMII) heavy chain to initiate apical constriction (AC) during *Xenopus* neural tube closure. Lmo7 is not required for actomyosin pulsatile activity or changes in the junction length in the intercalating cells during anteroposterior axis elongation. However, Lmo7 knockdown fails to stabilize actomyosin at the apical cortex at the onset of neural tube folding. Gain-of-function approach in gastrula ectoderm confirms a role for Lmo7 in actomyosin stabilization. Mechanistically, force-dependent dephosphorylation of Ser355 in the Lmo7 myosin binding domain enhances Lmo7 binding to NMII and increases NMII abundance at the apical cortex. Notably, initially homogeneous expression of Lmo7 in ectoderm cells progressively leads to apical domain heterogeneity that tightly correlates with Lmo7 levels, arguing for a positive feedback regulation between mechanical forces and Lmo7 activity. We propose that the force-dependent binding of Lmo7 to NMII stabilizes both proteins at the apical cortex, triggering enhanced apical constriction.

## Introduction

Apical constriction (AC), the reduction of apical domain size, is a fundamental cell shape change during epithelial morphogenesis (Perez-Vale and Peifer, 2020; Sawyer et al., 2010). One consequence of AC is cell ingression, during which AC cells dissolve apical junction and delaminate from epithelia, a process called epithelial-to-mesenchymal transition (EMT)(Baum et al., 2008; Francou et al., 2023; Ramkumar et al., 2016; Williams et al., 2012). Another consequence is epithelial folding, during which a group of cells coordinately increase actomyosin contractility and reduce their apical domains. Failure of coordinated AC results in severe congenital abnormalities, such as defects in neural tube closure (NTC) (Colas and Schoenwolf, 2001; Eiraku et al., 2011; Sherrard et al., 2010; Wallingford et al., 2013).

Actomyosin contractility generates mechanical forces required for AC and cell-cell rearrangement and is dynamically reorganized during epithelial morphogenesis. Actomyosin networks at the medioapical cortex undergo multiple cycles of contraction, stabilization and relaxation, generating actomyosin pulses. At the onset of AC, a “ratchet” stabilizes the transient contraction of actomyosin, reducing the size of apical domain and apical junctions (Martin and Goldstein, 2014; Martin et al., 2009; Miao and Blankenship, 2020). Many findings have emerged from studies in *Drosophila* embryos, and their importance has been established. Recent studies have reported similar actomyosin pulses in vertebrates that appear to be important for epithelial folding (Baldwin et al., 2023; Butler and Wallingford, 2018; Lardennois et al., 2019; Maitre et al., 2015; Shindo et al., 2019; Zhou et al., 2015). However, it remains unknown how medial actomyosin is stabilized in cells with oscillatory pulsation during epithelial folding.

Furthermore, coordinated AC in vertebrates is highly heterogeneous and asynchronous. Folding epithelia, such as the neuroectoderm, have cells with highly variable apical domain size, shape, and orientations (Baldwin et al., 2022; Christodoulou and Skourides, 2015; Colas and Schoenwolf, 2001; Schoenwolf and Smith, 1990; Suzuki et al., 2017), compared to AC in invertebrates which is regulated by specialized signaling pathways (Benton et al., 2019; Costa et al., 1994; Manning et al., 2013; Martin and Goldstein, 2014). At the onset of NT folding, apical domain heterogeneity is first apparent in the hinge regions of the neuroectoderm (Copp and Greene, 2010; Haigo et al., 2003; Matsuda et al., 2023; Sawyer et al., 2010). Isotropic AC cells increase actomyosin contractility and generate tensile force that pulls on their neighbors, in which actomyosin contractility is decreased along the anteroposterior (AP) axis, leading to the elongated apical domain (apical elongation, AE). This anisotropy in actomyosin contractility is important for preserving the length of the NT during its folding. The importance of anisotropic actomyosin contractions has been also reported in the neuroectoderm during early stages of neurulation, where the convergent extension (CE) elongates the neuroectoderm along the AP axis by the shrinkage of mediolaterally (ML)-oriented junctions and the elongation of AP-oriented junctions (Baldwin et al., 2022; Christodoulou and Skourides, 2022; Matsuda and Sokol, 2025). Although actomyosin contractility decrease has been considered a passive event, recent studies in *Drosophila* has identified regulatory proteins, such as PTEN, that facilitate junction elongation (Bardet et al., 2013; Ikawa et al., 2023; Uechi and Kuranaga, 2019; West et al., 2017), underscoring the importance of active AE in folding epithelia, while it is unknown whether their counterparts in vertebrates have similar functions.

Lmo7 is a multidomain scaffold protein located at apical junctions and the apical cortex (Beati et al., 2018; Du et al., 2019; Ooshio et al., 2004). Lmo7 overexpression (OE) induces coordinated AC in the *Xenopus* gastrula ectoderm, while Lmo7 knockdown (KD) disrupts NTC (Matsuda et al., 2022). Lmo7 binds and recruits non-muscle II (NMII) heavy chains to the apical cortex and junctions, increasing the formation of actomyosin networks (Matsuda et al., 2022).

Notably, modest overexpression of exogenous Lmo7 (low Lmo7 OE) induces the formation of AC and AE cells in an initially homogeneous cell population in the gastrula ectoderm (Matsuda et al., 2023). The increased apical domain variability in low Lmo7 OE resembles that in the folding neuroectoderm. We hypothesize that Lmo7 anisotropically promotes the reorganization of actomyosin network within the neuroectoderm, increasing the apical domain heterogeneity in the folding NT.

We first show that Lmo7 is not required for actomyosin pulsation at the apical cortex that remodels junctions in the neuroectoderm undergoing CE. At the onset of NT folding, however, the abundance of apical actomyosin failed to increase in Lmo7 KD cells, attenuating AC. Gain-of-function approach shows that Lmo7 has a specific activity on stabilizing pulsatile actomyosin at the apical cortex. A previous study reported Ser dephosphorylation in the NMII binding domain of Lmo7 in response to compressive force (Hashimoto et al., 2019). Our data suggest that the phosphorylation of the Ser and nearby Thr inhibits Lmo7 binding to NMII, AC and the generation of apical domain heterogeneity in the gastrula ectoderm. Based on these findings, we propose that Lmo7 is a force-responsive positive regulator of actomyosin and AC, necessary for neural plate folding.

## Results

### Lmo7 KD attenuates apical constriction, disrupting NTC

Lmo7 knockdown (KD) in the neuroectoderm disrupted NTC (Matsuda et al., 2022); however, the molecular and cellular mechanisms underlying this defect have not been studied. To better understand how Lmo7 regulates actomyosin or AC during NTC, we assessed the effects of Lmo7 KD on the dynamics of cells and junctions in the neuroectoderm. To compare the behaviors of cells and junctions in the same embryos, we injected Lmo7 morpholino (MO) on one side of the NE with RNA encoding lifeact-mNeon, and Lifeact-mScarlet RNA on the other side (Fig. 1A). In the control wild-type neuroectoderm, the apical domain size decreased rapidly at the onset of NT folding (magenta in Fig. 1B and Movie 1; Fig. 1D). In the Lmo7 KD NE, the apical domain remained significantly larger (green in Fig. 1B and Movie 1; Fig. 1D). We further confirmed the increased apical domain size through phalloidin staining (Fig. 1C). These findings confirm that Lmo7 is required for AC during NTC.

**Figure 1.**
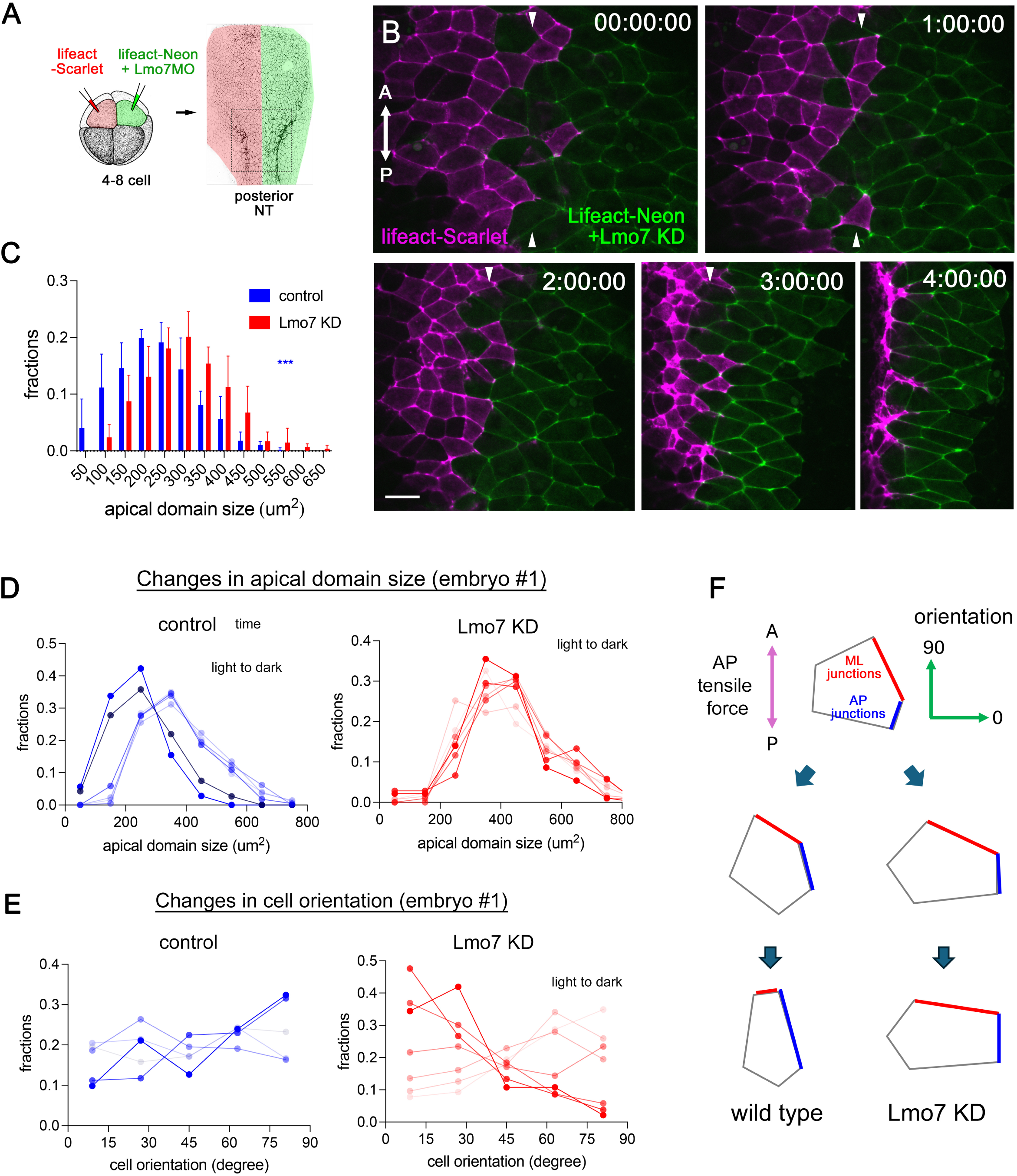
Lmo7 knockdown in the Xenopus neuroectoderm disrupts NTC. (A) Experimental scheme: Lmo7 morpholino (MO) was injected with lifeact-mNeon RNA into one dorsal animal blastomere at the 4-8 cell stage (green). Lifeact-mScarlet RNA, serving as a control, was injected into the other dorsal animal blastomere (magenta). Imaging was conducted on the superficial neuroectoderm (NE) of the posterior neural tube near the brain‒spinal cord border. (B) Selected images from Movie 1, showing control and Lmo7 knockdown (KD) cells in the NE undergoing NTC. Imaging started at stage 13, with 5 min intervals over a duration of 4 hours. Arrowheads indicate the presumptive midline. A: anterior; P: posterior. (C) Frequency distribution histogram of apical domain size at stage 15. The total number of cells from three independent embryos: n=620 (control) and n=312 (Lmo7 KD). (D, E) Representative changes in apical domain size (D) and cell orientation (E) in the NE. Frequency distribution histograms were based on time-lapse imaging data from one embryo, with five (control) and six (Lmo7 KD) time points. Darker dots represent NEs at later stages. Circular angles 0 and 90 in (E) correspond to the anteroposterior (AP) and mediolateral (ML) axes, respectively. The number of cells from each time point: 197, 215, 210, 220, 187, and 71 cells for control; 129, 135, 118, 111, 103, 105, and 93 cells for Lmo7 KD. Similar analysis was conducted on another independent embryo (Fig. S1). (F) Schematic representation of cell behaviors in the control wild type and Lmo7 KD NEs. In the wild type NEs, ML junctions (red) preferably shrink while AP junctions (blue) elongate, reorienting cells from ML to AP. In the Lmo7 KD NE, cells are preferably AP elongated during early neurula stages, while cells exhibit ML orientation in later stages. The Kolmogorov-Smirnov test was used to assess distribution differences between the control and Lmo7 KD NEs in (C), and between the first and last frames in (D) and (E). ***p<0.001. Scale bar: 20 μm (B).

Another change observed in the Lmo7 KD NE was cell orientation. The extrinsic and intrinsic forces highly control the behaviors of cells and junctions in the NE. In the early neurula, CE of the underlying mesoderm was proposed to pull the NE along the AP axis, leading to the preferential shrinking of ML-oriented junctions and the elongation of AP-oriented junctions (Baldwin et al., 2022; Christodoulou and Skourides, 2022; Matsuda and Sokol, 2025).

Subsequently, increased actomyosin contractility initiates isotropic AC in a subset of cells first at neural hinges, associated with the AP elongation of neighboring cells. These orientation-based behaviors progressively increase the AP-oriented cell population in the control wild type NE (lighter to darker blue in Fig. 1E). In the Lmo7 KD NE, cells preferentially AP-oriented in early stage neurula became ML-oriented in later stages (lighter to darker red in Fig. 1E). These results suggest that Lmo7 is required for the orientation-based behaviors of cells and junctions in the folding NE (Fig. 1F).

### Lmo7 KD does not significantly alter PCP or the orientation of cells and cell junctions in the neuroectoderm

The planar cell polarity (PCP) signaling pathway coordinates cell behaviors in the plane of a tissue, playing an important role in orienting cells and junctions during epithelial folding, including NTC (Ciruna et al., 2006; Ybot-Gonzalez et al., 2007). Recently, we reported that Prickle 2 (Pk2), a core component of PCP signaling, is required for the orientation-based junction behaviors during NTC (Matsuda and Sokol, 2025). In the Pk2 KD NE, AP and ML junctions did not preferentially elongate or shrink. Pk2 KD cells remained ML-oriented later in neurulation, which resembles Lmo7 KD cells in the NE (Fig. 1B, E, F). Thus, we asked whether Lmo7 controls the behaviors of cells or junctions through examining Pk2 localization.

First, we evaluated the establishment of PCP in the Lmo7 KD NE. The hallmark of PCP is the planar polarized distribution of two core PCP complexes at the opposite sides of cells (Humphries and Mlodzik, 2018; Peng and Axelrod, 2012). In the NE, GFP-Pk2 forms a complex with Vangl2 and preferentially localizes at the anterior end of cell cortex (arrowheads in the control NE in Fig. S2A)(Butler and Wallingford, 2018; Matsuda and Sokol, 2025). Compared to the loss of GFP-Pk2 polarization in Vangl2 KD, GFP-Pk2 preferentially localized at the anterior junctions in the Lmo7 KD NE (arrowheads in Fig. S2A). Next, we assessed the orientation-dependent behaviors of cells and junctions in Lmo7 KD cells. AP junctions were found to preferentially elongate, while ML junctions preferably shrank, as observed in the control wild-type cells (Fig. 2D). These results suggest that the loss of Pk2 polarization is unlikely the primary cause of NTC defects in the Lmo7 KD embryos.

**Figure 2.**
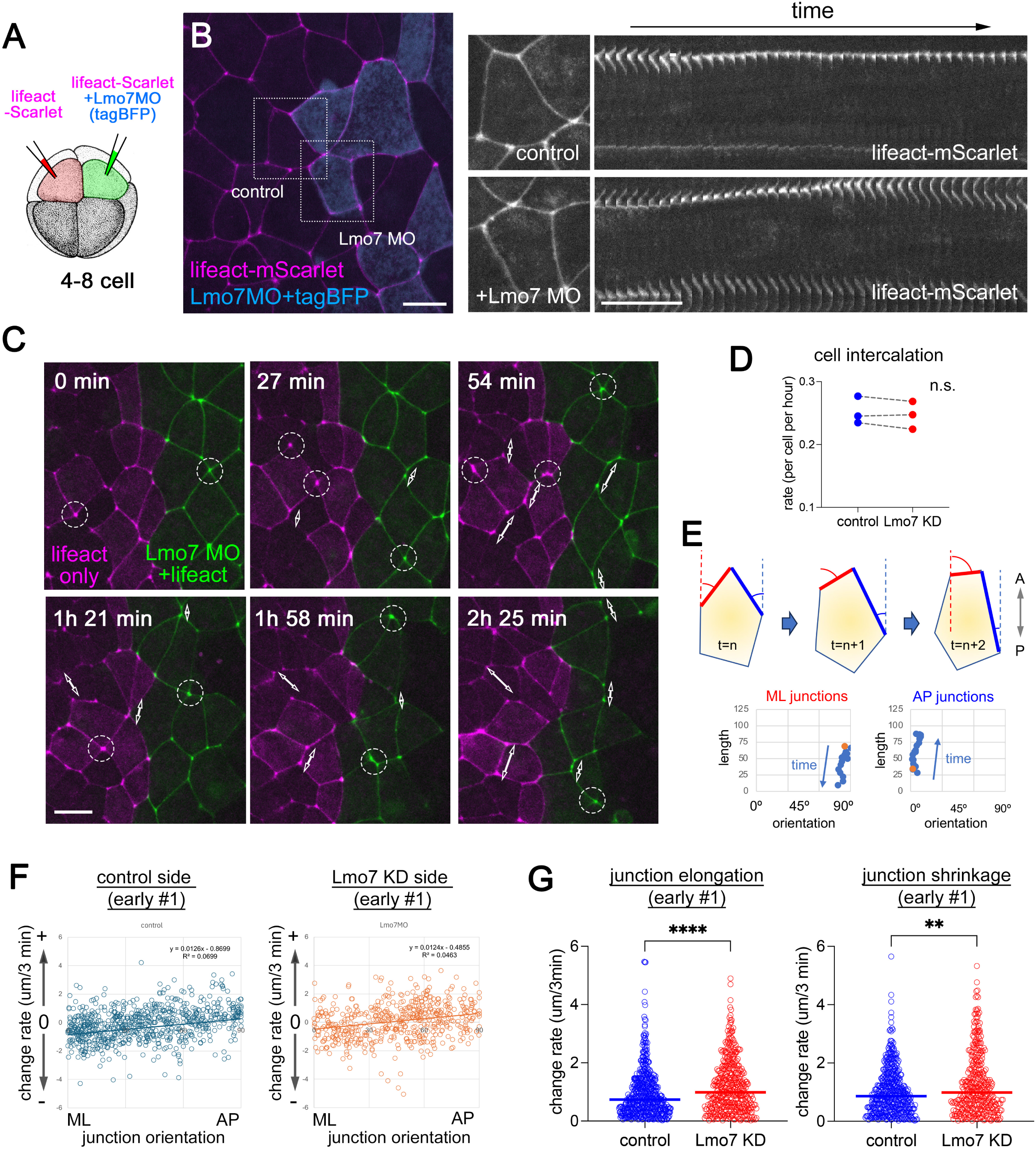
The effects of Lmo7 knockdown on cells and junctions in the neuroectoderm undergoing AP-oriented convergent extension. (A) Experimental scheme: lifeact-mScarlet was expressed in both sides of the NEs, while Lmo7 MO was coinjected into one side along with tagBFP as a tracer. Time-lapse imaging began at stage 14 in the posterior NE near the midline where the control and Lmo7 KD cells were in the same imaging field. Two square areas in (A) are enlarged in (B) for control and (C) for Lmo7 KD. (B) Selected frames and representative kymograph from Movie 2. Imaging duration was 19 min and 30 s with 30-s intervals. Pulsatile F-actin is visible in both control and Lmo7 KD NEs. (C) Selected images from Movie 3, showing the effects of Lmo7 KD on junction length changes and cell intercalation. Lifeact-mScarlet was expressed on one side, and lifeact-mNeon with Lmo7 MO on the other side, as illustrated in Fig. 1A. Imaging started at stage 14, with intervals of 4 minutes over a duration of 2 hrs and 56 min. Circles indicate T1 junctions. Double-headed arrows indicate new junctions that elongate following T1 resolution. Arrowheads: the presumptive midline. A: anterior; P: posterior. (D) The frequency of cell intercalation in the posterior neuroectoderm during the early neurula stage. Paired dots represent the frequency of cell intercalation for the control and Lmo7 KD sides within the same embryos, with data from three embryos. The total cell counts from individual embryos are as follows: 65, 52, and 115 cells for control, and 67, 97, and 52 cells for Lmo7 KD. (E) Schematic representing how the orientation and length changes of junctions are quantified through frame-by-frame analysis. Shrinking ML junctions (red) and elongating AP junctions. (F) Two-dimensional plots represent the orientation (x-axis) and length change (y-axis) of individual junctions over the first 2 hrs and 30 min of time-lapse imaging in Movie 1. Circular angles of 0 and 90 correspond to ML and AP axes, respectively. Positive and negative junction length values indicate elongation and shrinkage. (G) The rates of junction elongation and shrinkage, utilizing the same dataset as in (F). Bold and thin lines represent the medians and quartiles, respectively. The total number of junction changes analyzed includes 546 elongations and 501 shrinkages for the control, and 441 elongations and 387 shrinkages for Lmo7 KD. An independent analysis was performed on another embryo (Fig. S2B and C). Statistical significance was assessed using a paired Student’s t-test for means of cell intercalation rates in (E), and the Mann-Whitney test for mean ranks of junction elongation and shrinkage in (G). **p<0.01, ****p<0.0001, n.s. not significant. Scale bars in B and C: 20 μm.

### Lmo7 is not required for actomyosin pulsation at the apical cortex or for the efficient remodeling of junctions during elongation or shrinkage

Previous studies in *Drosophila* have shown that the oscillatory constriction and relaxation of the actomyosin network at the apical cortex (Martin et al., 2009). A “ratchet” mechanism has been proposed to stabilize actomyosin pulses, gradually reducing the size of the apical domain (Martin and Goldstein, 2014; Martin et al., 2009; Miao and Blankenship, 2020). Recent studies in *Xenopus* embryos showed that medioapical actomyosin pulsates in the NE and becomes more abundant in AC cells during NTC (Baldwin et al., 2022; Christodoulou and Skourides, 2022). The same study also reported the absence of medioapical actomyosin increase in the NE depleted of Shroom3, a well-established AC inducer (Das et al., 2014; Haigo et al., 2003; Hildebrand, 2005; Hildebrand and Soriano, 1999). Indeed, loss of medioapical actomyosin increase appears to correlate with AC detects in *Xenopus* embryos (Baldwin et al., 2023). Since Lmo7 OE in the gastrula ectoderm enhanced actomyosin network both at apical junctions and the apical cortex (Matsuda et al., 2022), we next examined the effects of Lmo7 KD on actomyosin dynamics in the folding NE. Since the posterior NE in *Xenopus* embryos have two phases of cell behaviors (Baldwin et al., 2022; Christodoulou and Skourides, 2022), we separately analyzed the effects of Lmo7 KD in early and late stage neurula.

Time-lapse imaging was first conducted to assess the effects of Lmo7 KD on actomyosin dynamics in the early stage neurula during convergent extension (Fig. 2A, B). Actomyosin networks labeled by Lifeact-mScarlet were pulsatile, constantly relocating within the medioapical cortex in the control wild-type NE (Fig. 2C, Movie 2), consistent with previous reports (Baldwin et al., 2022). Actomyosin pulses were present in Lmo7 KD cells in the same embryos (Fig. 2D, Movie 2). The spatial pattern and frequency of actomyosin pulses were also comparable between wild-type and Lmo7 KD cells, although we acknowledge that our imaging may be limited by sensitivity and time resolution. Consistent with the unchanged actomyosin pulses, Lmo7 KD did not significantly alter the direction or frequency of cell intercalation (Fig. 2E, F, Movie 3), or the orientation-based preference of junction length change (Fig. 2G, Movie 3). The rates of junction elongation and shrinkage was slightly increased (Fig. 2J), while statistical significance was marginal in some embryos (Fig. S2B, C). Of note, Pk2 KD generally attenuated junction remodeling and reduced the rates of junction elongation and shrinkage in the NE (Matsuda and Sokol, 2025), which was opposite to the effects of Lmo7 KD on junction remodeling. These results suggest that Lmo7 is not required for actomyosin pulsation or junction remodeling for proceeding cell intercalation during CE.

### Lmo7 KD fails to stabilize actomyosin at the medioapical cortex in the folding NE

To test the effects of Lmo7 KD on actomyosin dynamics during NT folding, we utilized the NE coexpressing Lifeact-mScarlet and Sf9-mNeon, live probes for F-actin and NMIIA (Vielemeyer et al., 2010) (Fig. 3A). First, both Lifeact and Sf9 increased the abundance in apically constricting cells in the wild type control NE (Fig. 3C), recapitulating the effects of Lmo7 OE on endogenous F-actin. Time-lapse imaging showed the stabilization of medioapical actomyosin pulses in the control side of the NE (left sides in Fig. 3B, C, Movie 4). In contrast, actomyosin continued to pulsate in Lmo7 KD cells and their abundance remained low at the apical cortex (right side in Fig. 3B, C, Movie 4). The apical domain of Lmo7 KD cells also remained large, consistent with the reduced rate of junction shrinkage (Fig. 3F). Phalloidin staining confirmed medial F-actin in the Lmo7 KD neuroectoderm as compared to the controls (Fig. S3B).

**Figure 3.**
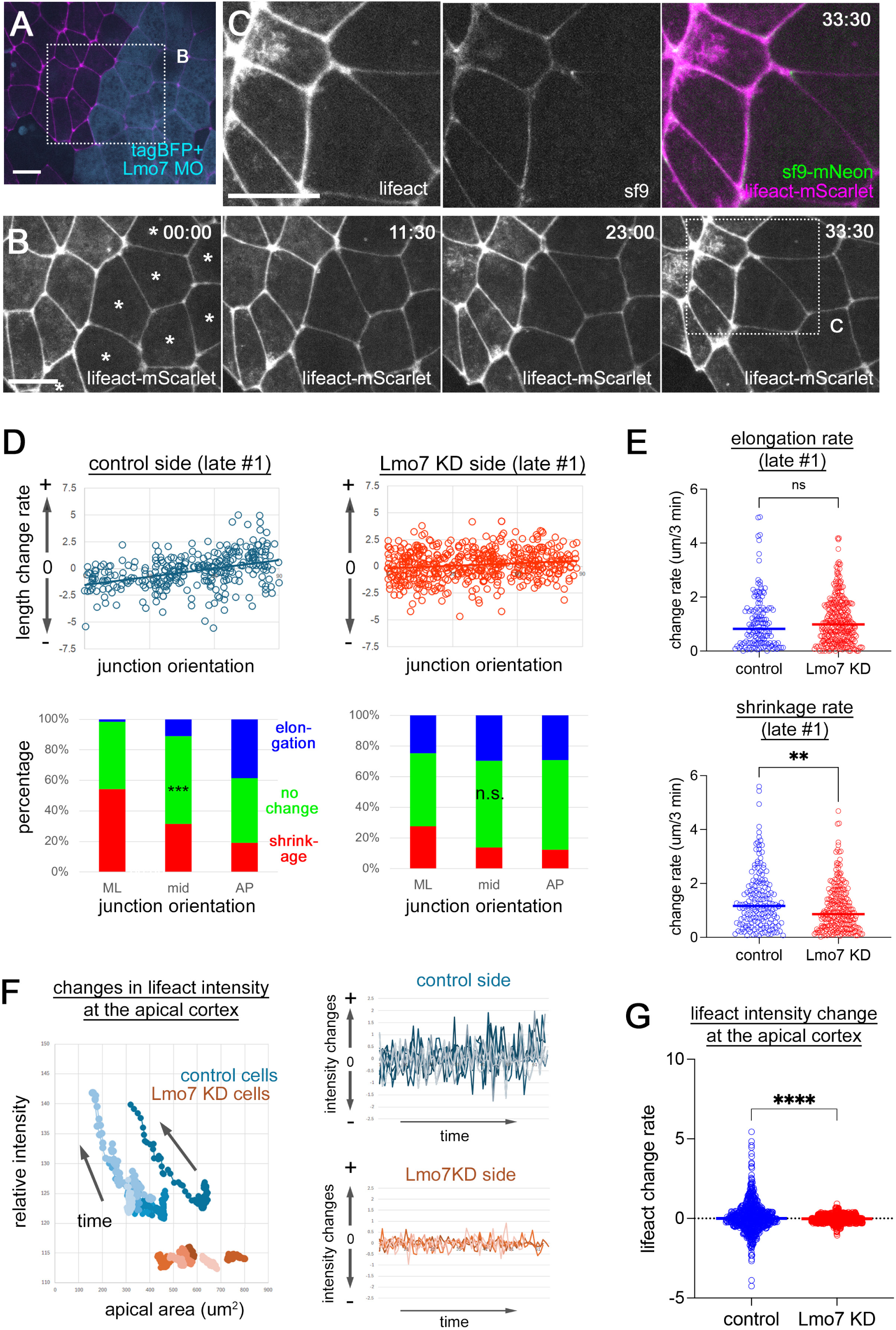
Lmo7 knockdown suppresses actomyosin increases at the apical cortex in folding Nes. (A) A representative image of the NE before time-lapse imaging. Both sides of the NE express lifeact-mScarlet and sf9-mNeon. Lmo7 MO was injected into only one side, with tagBFP RNA serving as a tracer. The square area is enlarged in panel B. Arrowheads indicate the presumed midline. A: anterior; P: posterior. (B) Selected images from Video 4, showing only lifeact-mScarlet channel only. Time-lapse imaging began at stage 16, with intervals of 30 s for a duration of 34 min. The square area is enlarged in panel C. Asterisks indicate Lmo7 KD cells. (C) Merged images of lifeact-mScarlet and sf9-mNeon in the control (left) and Lmo7 KD (right) NEs. Arrowheads indicate actomyosin at the medioapical cortex. (D) Two-dimensional plots illustrating the orientation (x-axis) and length changes (y-axis) of individual junctions. Junctions were randomly selected from one time-lapse imaging, with control cells on one side and Lmo7 KD cells on the other. Circular angles of 0 and 90 correspond to ML and AP axes, respectively. Positive and negative values in length change indicate elongation and shrinkage of junctions, respectively. Junction length changes were categorized as elongation (blue, ≥1 mm/3 min), shrinkage (red, ≤-1 mm/3 min), and no change (green,-1 mm < change <1 mm/3 min). Junctions were classified into APs (≤30° from the AP axis), MLs (≤30° from the ML axis), and mid (30-45° from either the AP or ML axes). A two-sample Kolmogorov-Smirnov test assessed the distribution differences between AP and ML junctions using the original unmetrical data. The total number of junction length changes was as follows: n=142 (control elongation), n=168 (control shrinkage), n=298 (Lmo7 KD elongation), and n=237 (Lmo7 KD shrinkage). (E) Overall rates of junction elongation (top) and shrinkage (bottom) were derived from the same data in panel D. Bold and thin lines represent the medians and quartiles, respectively. The Mann-Whitney test compared the mean ranks. (F) Quantification of lifeact-mScarlet at the apical cortex. Left: A two-dimensional plot (left) with apical domain size on the x-axis and fluorescence intensity of lifeact-mScarlet on the y-axis. Dots and lines connecting dots represent five independent control (blue) and Lmo7 KD (red) cells which were randomly selected and tracked in Movie 4. Right: one-dimensional plots with time on the x-axis and intensity changes in lifeact-mScarlet on the y-axis. (G) Fluorescence intensity was quantified in late stage neurula where AC was visible on the control side of the NEs. Three embryos per group. Total number of cells: n=755 (control elongation) and n=1114 (Lmo7 KD). **p<0.01, ****p<0.0001, n.s., not significant. Scale bars in A, B and C: 20 μm.

Notably, as the control side of the NE progressively increased AC cell population, they appeared to pull adjacent Lmo7 KD cells medially (Figs. 1B, 3B, Movies 1, 4). AP and ML junctions elongated and shrank with similar frequencies in the Lmo7 KD NE (Fig. 3D). This supports the idea that the Lmo7 KD NE cells in late neurulas are affected by both AP- and ML-oriented tensile forces, independently of their orientation. Taken together, these results suggest that Lmo7 KD fails to stabilize actomyosin pulses in folding NT.

### Lmo7 overexpression stabilizes apical actomyosin in gastrula ectoderm

In general, AC defects are associated with loss of actomyosin increase at the apical cortex in *Xenopus* embryos (Baldwin et al., 2022; Matsuda et al., 2023). Therefore, loss of medial actomyosin accumulation in the Lmo7 KD NE does not indicate a specific causal role for Lmo7 in stabilizing actomyosin at the apical cortex. To test the instructive role of Lmo7 in actomyosin stabilization, gain-of-function (GOF) approaches were taken. We utilized the embryonic ectoderm system in gastrula-stage embryos due to its simplicity and its suitability for high-resolution imaging and well-established apical junctions. Using this system, we previously demonstrated that Lmo7 overexpression (OE) promotes the formation of actomyosin networks at both apical junctions and the medioapical cortex (Matsuda et al., 2022).

Live probes for F-actin and NMII were coexpressed with Flag-Lmo7 (high, 500 pg RNA) in the gastrula ectoderm (Fig. 4A). High level Lmo7 OE increased the formation of actomyosin networks at both junctions and the apical cortex (middle panels in Fig. 4B), recapitulating the effects of Lmo7 OE on endogenous F-actin and NMII (Matsuda et al., 2022). In time-lapse imaging of wild-type ectoderm, apical actomyosin pulses were short-lived and formed at random locations (control in Fig. 4C, D, and Movie 5). In the Lmo7 OE ectoderm, actomyosin pulses stayed at the same location in the apical cortex for a prolonged duration (Lmo7 KD in Fig. 4C, D and Movie 5). The increased stabilization of apical actomyosin in the Lmo7 OE ectoderm contrasts with its loss in the Lmo7 KD NE (Fig. 3).

**Figure 4.**
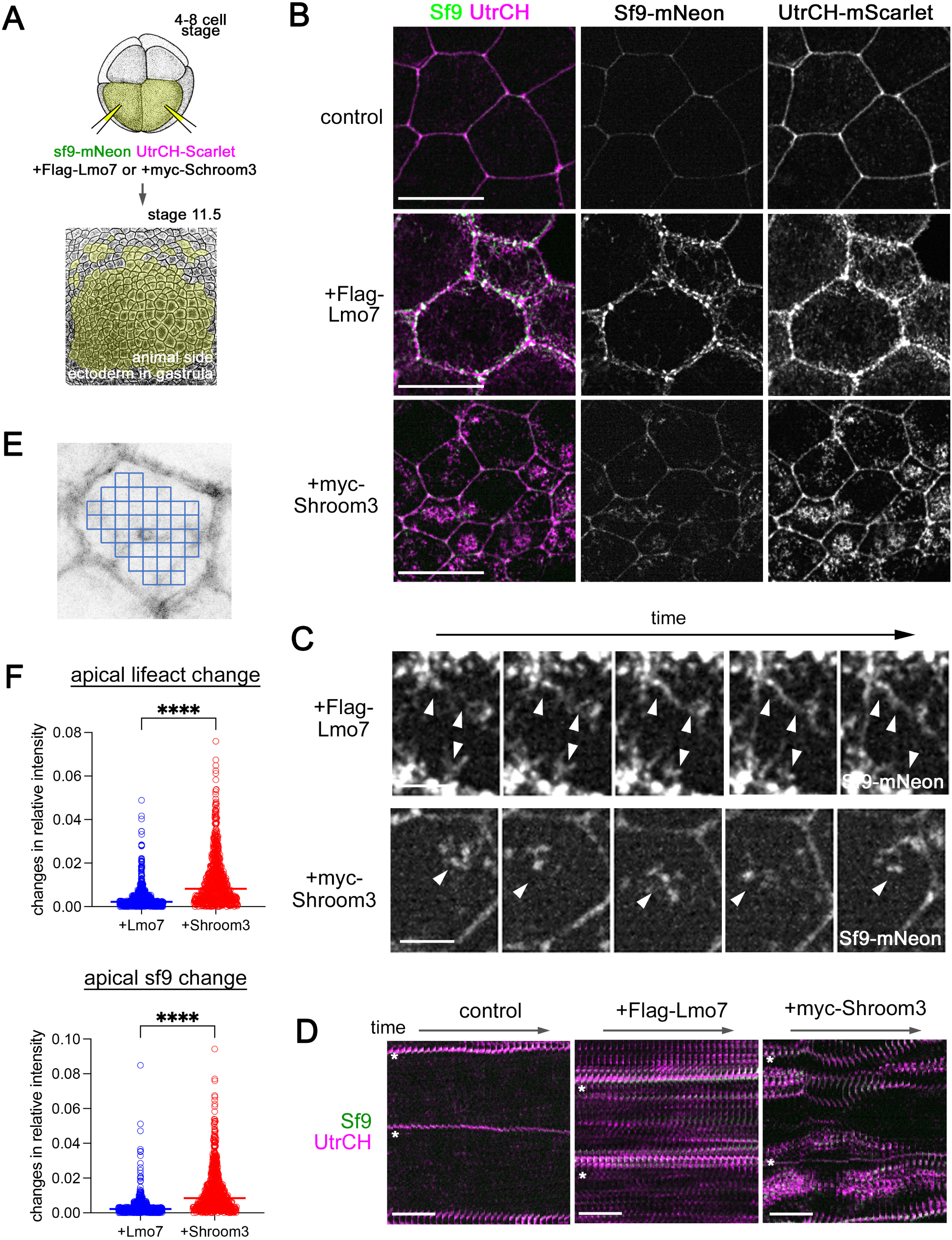
Lmo7 OE stabilizes apical actomyosin in the gastrula ectoderm. (A) Experimental scheme: Flag-Lmo7 or myc-Shroom3 RNA was co-injected with UtrCH-mScarlet and sf9-mNeon into two ventral blastomeres at the 4 or 8 cell stage. Time-lapse imaging of the animal superficial ectoderm began at stage 11. (B) Representative still image of the gastrula ectoderm expressing UtrCH-mScarlet and sf9-mNeon only, with Flag-Lmo7 or with myc-Shroom3. (C) Selected frames from Movie 5, showing actomyosin dynamics. Sf9-mNeon channel only is shown. Imaging started at stage 11, with 30-s intervals over a duration of 15 min. (D) Representative kymographs from Movie 5. (E) Schematic illustrating actomyosin dynamics are quantified at the apical cortex. The apical domain was divided into 2 by 2 micrometer squares, and the fluorescence intensity of sf9-mNeon and lifeact-mScarlet was quantified in squares that did not include any apical junctions. (F) Changes in intensity of sf9-mNeon and lifeact-mScarlet at the apical cortex. The total number of squares analyzed: 716 squares (+Flag-Lmo7) and 493 squares (+myc-Shroom3) from >10 cells per conditions. The Mann-Whitney test was used to compare mean ranks. ****p<0.0001. Scale bars: 20 μm (B), 5 μm (C), 10 μm (D).

Notably, the immobilization of actomyosin pulses at the apical cortex was specific to Lmo7 OE. Shroom3 triggers actomyosin contractility and AC via the Rho-Rock-NMII pathway during NTC (Das et al., 2014; Haigo et al., 2003; Hildebrand, 2005; Hildebrand and Soriano, 1999). Failure of AC in the Shroom3 KD NE was accompanied by the lack of medial actomyosin increase in *Xenopus* embryos (Baldwin et al., 2022). In our analysis, Shroom3 OE enhanced the formation of actomyosin networks at the apical cortex, as indicated by the increased abundance of Lifeact and Sf9 in the gastrula ectoderm (+myc-Shroom3 in Fig. 4B). Despite this, actomyosin in the Shroom3 OE ectoderm were highly pulsatile and dynamically changed location within the apical cortex (green arrows in Fig. 4C, D, Movie 5). We quantified the changes of Lifeact or Sf9 intensity in small squares at the apical cortex (Fig. 4E). Actomyosin dynamics in the Shroom3 OE was higher than those in the Lmo7 OE ectoderm (Fig. 4F). A failure of PCP establishment by Pk2 or Vangl2 KD disrupted NTC, which was also associated with a loss of AC and medial actomyosin accumulation (Baldwin et al., 2022; Christodoulou and Skourides, 2022; Matsuda et al., 2023; Matsuda and Sokol, 2025). However, Pk2 OE did not increase or stabilize medial actomyosin (Matsuda and Sokol, 2025). Taken together, these results suggest that Lmo7 has a direct role in stabilizing medial actomyosin during AC.

### Positive regulatory feedback between Lmo7 and actomyosin

A previous study of the *Xenopus* gastrula reported that compressive force posttranslationally increased Lmo7 stability and this effect correlated with the decreased phosphorylation of the conserved Ser355 and an adjacent Thr in the NMII binding domain (NMBD) (Hashimoto et al., 2019). We reported that low-level Lmo7 OE progressively increased the heterogeneity of apical domain size, forming cells with small apical domains adjacent to those with larger ones (Matsuda et al., 2023). We also noticed a reversed correlation between apical domain size and Lmo7 abundance at the apical cortex. This suggests a mutual regulation between Lmo7 and actomyosin force.

To test whether Lmo7 is force-sensitive, a low dose of GFP-Lmo7 (100 pg RNA) was injected into two ventral blastomeres, along with membrane-tethered myrRFP or myrBFP to normalize the amount of incorporated RNA in individual cells (Fig. 5B). Both GFP-Lmo7 and myrRFP or tagBFP signals were relatively uniform among cells at the beginning of time-lapse imaging (00:00 or t=0 in Fig. 5C, E, F, G, Movie 6). We tracked apical domain size and GFP-Lmo7 intensity at the apical cortex in individual cells (Fig. 5D). By the end of time-lapse imaging, GFP-Lmo7 intensity at the apical cortex became heterogeneous, while myrRFP/BFP signals remained uniform (00:00 or t=27 in Fig. 5C, E, F, Movie 6). These results suggest that the increased variability of GFP-Lmo7 among cells is post-translationally regulated and not due to variability in the amount of injected RNA. Furthermore, GFP-Lmo7 intensity was greater in cells with constricting apical domain (Fig. 5C, D, Movie 6). These results suggest that a progressive increase in Lmo7 abundance enhances actomyosin contractility, leading to AC in these cells. In contrast, cells with low Lmo7 abundance maintain both low levels of Lmo7 and a larger apical domain.

**Figure 5.**
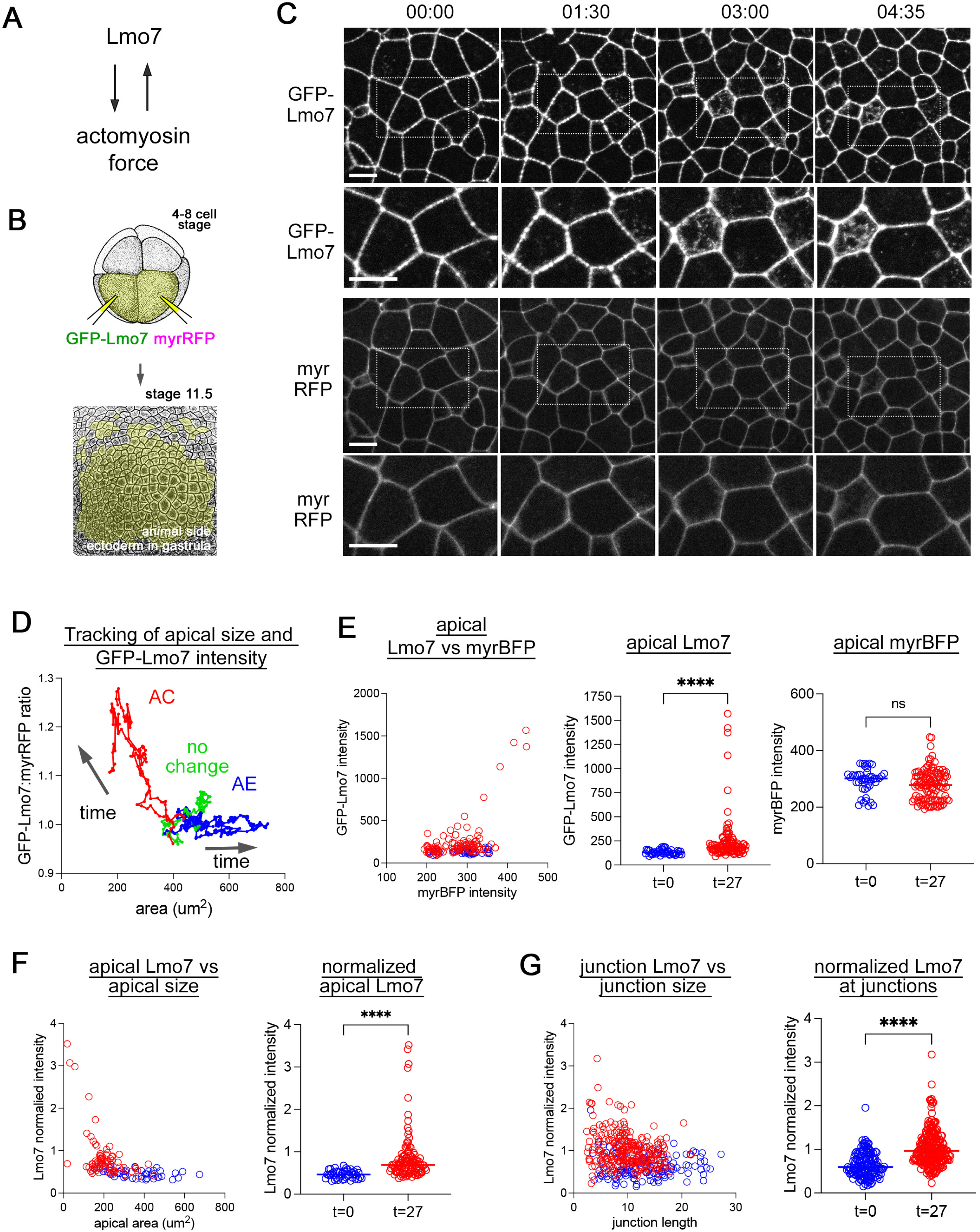
Lmo7 OE increases apical domain heterogeneity in the gastrula ectoderm. (A) Schematic illustrating the feedback regulation between Lmo7 and actomyosin force. (B) Schematic of the experimental setup: GFP-Lmo7 RNA was co-injected with myrRFP or myr-tagBFP-HA RNA into two ventral blastomeres at the 4-8 cell stage. (C) Selected images from Movie 6. The square areas are enlarged in the lower panels. Time-lapse imaging started at stage 11, with 5-minute intervals over a duration of 4 hours and 35 minutes. (D) A two-dimensional plot with apical domain size on the x-axis and normalized fluorescent GFP-Lmo7 intensity in the apical domain on the y-axis. Dots and lines represent individual cell tracking. Blue dots show 4 cells that increased apical domain size by more than 20%. Red dots show 3 cells that decreased apical domain size by more than 20%. Green dots show 3 cells with less than 20% change. The intensity of GFP-Lmo7 was normalized by the intensity of co-injected myrRFP. (E-G) Quantification of progressive GFP-Lmo7 accumulation at the apical cortex (E, F) and junctions (G). The abundance of GFP-Lmo7 in the apical domain at the first frame (blue) was compared to that at the last frame (red) of the time-lapse imaging. Left in (E): A representative two-dimensional plot with relative myrBFP intensity on the x-axis and relative GFP-Lmo7 intensity on the y-axis in the apical domain. Middle in (E): A one-dimensional plot showing relative GFP-Lmo7 abundance and variability increase. Right in (E): A one-dimensional plot showing no change in relative myrBFP intensity. Left in (F): A representative two-dimensional plot with apical domain size on the x-axis and normalized GFP-Lmo7 intensity in the apical domain on the y-axis. Right in (F): A one-dimensional plot showing the normalized intensity of GFP-Lmo7 abundance at the apical cortex. Left in (G): A representative two-dimensional plot with junction length on the x-axis and the normalized GFP-Lmo7 intensity at junctions on the y-axis. Right in (G): A one-dimensional plot comparing the normalized GFP-Lmo7 intensity at individual junctions. Bold and thin lines represent the medians and quartiles, respectively. A total of 45 cells and 144 junctions (t=0), and 111 cells and 252 junctions (t=27) from a single embryo, were analyzed in E-G. The same analysis was conducted on another embryo (Fig. S3). The Mann-Whitney U test was applied to compare mean ranks. ****p<0.0001. n.s., not significant. Scale bars: 20 μm. 355S 357T 358S

### Phosphomimetic mutation in Ser and Thr in the NMBD suppresses Lmo7 interaction with NMII and apical domain heterogeneity by Lmo7

The S355 residue in the Lmo7 NMBD (Fig. 6A) is dephosphorylated in response to a compressive force (Hashimoto et al., 2019). To examine the effects of S355 phosphorylation on Lmo7 binding to NMII, we substituted S355 with phosphomimetic and nonphosphorylatable residues. In a co-immunoprecipitation (IP) assay in HEK293 cells, the S355D phosphomimetic mutation reduced the amount of NMII heavy chains coimmunoprecipitate by full-length Lmo7 (Fig. 6B, C). Lmo7 binding to NMII heavy chains was further reduced by additional phosphomimetic mutations on the evolutionarily conserved T357 and S357(S3D in Fig. 6A-C). Consistently, Lmo7(S3D) mutation also suppressed Lmo7-mediated NMIIA increase at junctions (Fig. 6C-E). Lmo7(S3D) also suppressed the apical accumulation of pigment granule and the formation of epithelial indentation (Fig. S5), indicators of coordinated reduction of apical domain in the *Xenopus* gastrula ectoderm (Haigo et al., 2003; Hildebrand, 2005). These results suggest that Lmo7 activity is positively regulated by force via the dephosphorylation of Ser355 and the adjacent residues in the NMBD.

**Figure 6.**
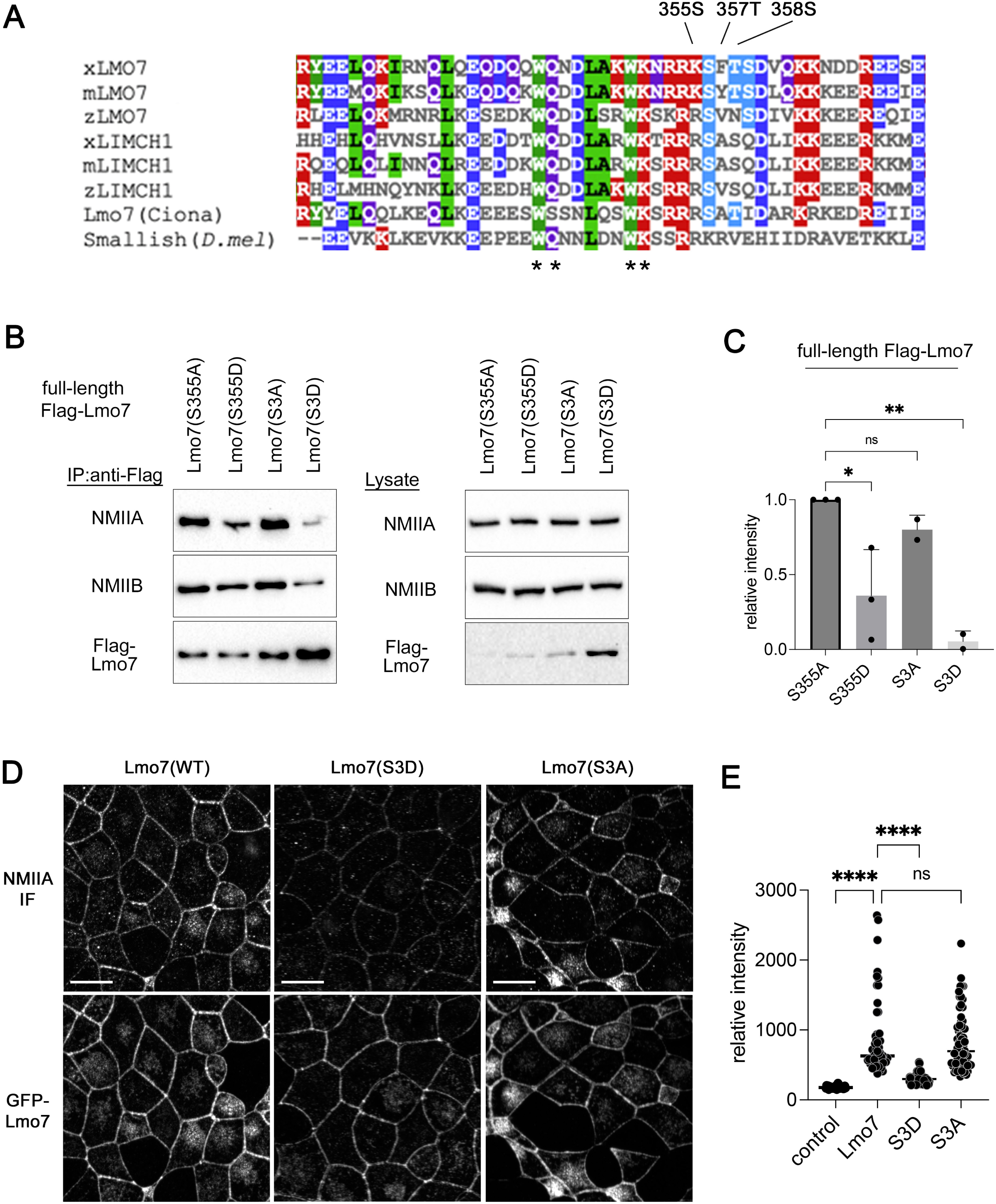
Phosphomimetic mutations in the NMII binding domain of Lmo7 suppress the co-IP and the apical enrichment of NMII by Lmo7. (A) Alignment of the NMII binding domain (NMBD) in Lmo7 and Limch1 in vertebrates, and Smallish in *Drosophila*. Conserved amino acids are color-coded: acidic (blue), basic (red), Ser or Thr (light blue), and hydrophobic (green). Asterisks indicate the conserved WQ-WK motif, which was substituted with Ala in the Lmo7(4A) mutant. (B) Co-immunoprecipitation assay conducted in HEK293T cells. Full-length Flag-Lmo7, either with phosphomimetic or non-phosphorylatable mutations in the NMBD, was immunoprecipitated using an anti-Flag antibody. The presence of endogenous NMIIA and NMIIB in Flag-Lmo7 IP products was evaluated by immunoblotting (IB). (C) The efficacy of NMIIA co-IP by Flag-Lmo7 was quantified from three independent experiments. Note that since the levels of Lmo7 mutant proteins in cell lysates are variable and higher than those of wild-type Lmo7, the band intensities of anti-NMIIA IB were normalized by those of anti-Flag IB and then relative to Lmo7(S3A). (D) Representative images showing NMIIA distribution in the gastrula ectoderm expressing GFP-tagged wild-type, S3D, and S3A mutant forms of Lmo7 at stage 11. The subcellular distribution of endogenous NMIIA was examined using anti-NMIIA immunofluorescence (IF). (E) Quantification of anti-NMIIA IF. The number of cells: n=30 (no Lmo7 control), n=54 (+Lmo7 (wild type)), n=64 (Lmo7(S3D)), and n=68 (Lmo7(S3A)). Three embryos per condition. One-way ANOVA was used to compare mean ranks, with Bonferroni correction for multiple pairwise comparisons in C and E. *p<0.05, **p<0.01, ****p<0.0001. n.s., not significant. Scale bars: 20 μm.

Furthermore, the Lmo7 with the S3D mutations exhibited reduced heterogeneity of apical domain size in the gastrula ectoderm (Fig. 7A-D, Movie 7). Lmo7 accumulation at apical junctions in the GFP-Lmo7(S3D) ectoderm was also more uniform than those in GFP-Lmo7 ectoderm (Fig. 7A, B, E, F, Movie 7). Notably, neither Lmo7(S355A) nor Lmo7(S3A) reduced Lmo7 binding to NMII (Fig. 6B, C) or the recruitment of NMII to the apical domain or junctions (Fig. 7D, E). They increased the accumulation in the apical cortex in apically constricting cells (Fig. 6D, E) and did not reduce apical domain heterogeneity and Lmo7 abundance at the apical cortex (Fig. 7A, B). Taken together, these results suggest that the formation of apical domain heterogeneity requires actomyosin contractility increase mediated by Lmo7, which is dependent on the dephosphorylation of Ser and Thr in the NMBD,

**Figure 7.**
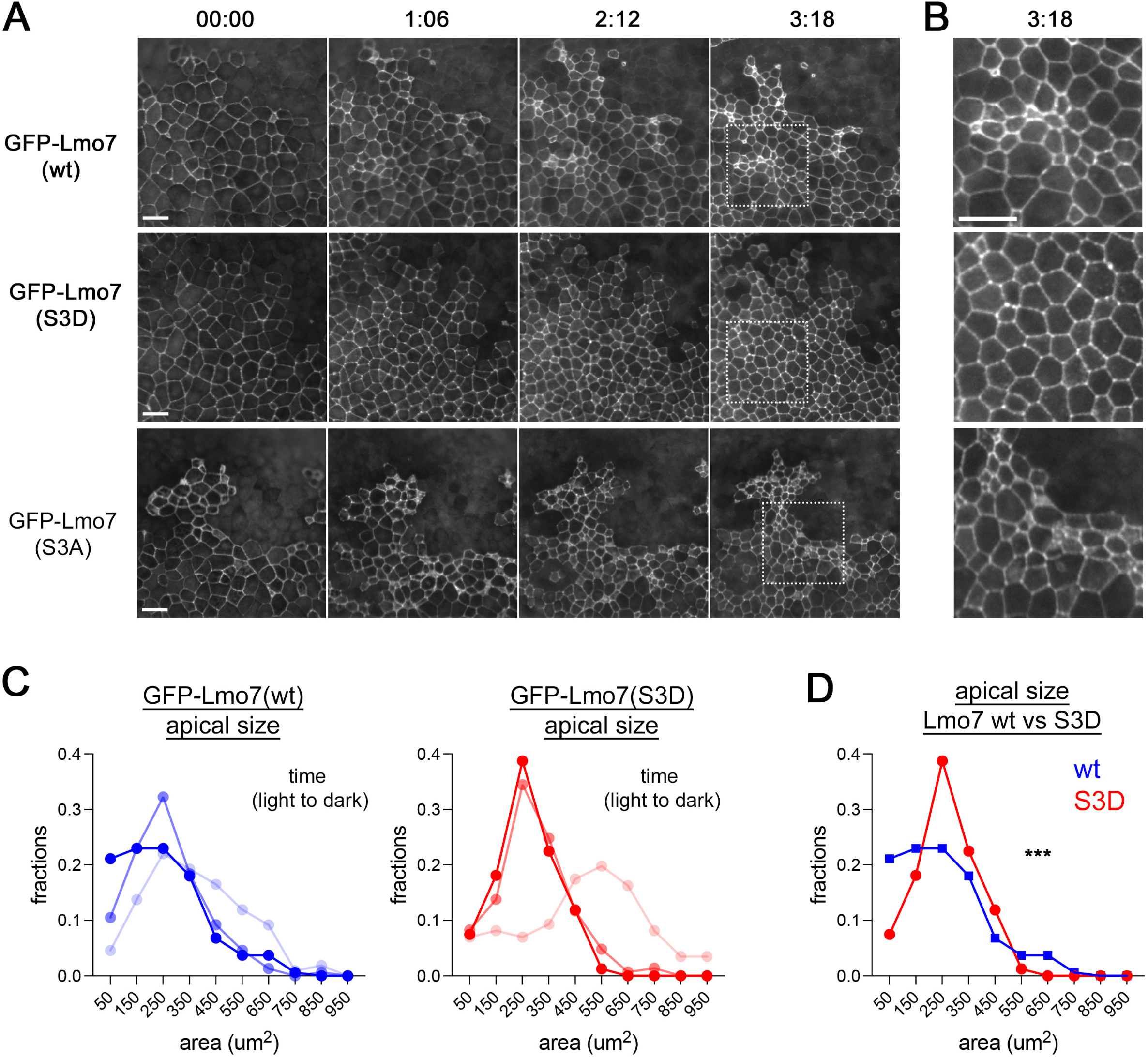
Phosphomimetic mutations of Ser and Thr suppress Lmo7 heterogeneity in the gastrula ectoderm. (A, B) Selected images from Movie 7, which shows gastrula ectoderms expressing low levels of GFP-Lmo7(wt), Lmo7(S3D), or Lmo7(S3A) from Movie 7. The square areas in A are enlarged in the same panel. Time-lapse imaging began at stage 11 with 6-min intervals over a duration of 3 hr 18 min. (C, D) Quantification of apical domain size heterogeneity from low-level OE of GFP-Lmo7 in blue and GFP-Lmo7(S3D) in red. Frequency distribution histograms in C illustrate the apical domain sizes of individual cells at three-time frames, with lighter colors representing earlier time points. Note that cell proliferation is highly active in the early-stage gastrula, leading to a reduction in apical domain size. The heterogeneity of apical domain size at the last time frame is presented in D. The total number of cells at each time point is: 109, 152, and 161 (wild type); 87, 145, and 160 (S3D). The two-sample Kolmogorov-Smirnov test was utilized to compare distributions in D. ***p<0.001. Scale bars: 40 μm.

### Conformational change in Lmo7 may be required for its effect on actomyosin network

We explored how phosphorylation in the NMBD controls Lmo7 activity on actomyosin.

One possibility is that phosphorylation in the NMBD may alter the conformation of Lmo7, allowing the binding interface of NMBD to be more accessible for NMII. Supporting this, S355D or S3D mutations in *full-length Lmo7* did reduce NMII co-IP by Flag-Lmo7 (Fig. 5A-C); however, the same mutations in the *NMBD fragment* did not significantly reduce NMII co-IP by Flag-NMBD (Fig. 8A, B). In contrast, 4A substitutions in the conserved WQ-WK sequence significantly reduced the efficacy of NMII co-IP by Flag-Lmo7 both in the full-length Lmo7 (Matsuda et al., 2022) and in the NMBD fragment (Fig. 8A, B). These results suggest that the 4A mutation directly alters the interface of the NMII binding site in the NMBD and abolishes NMII binding to the NMBD. Conversely, the phosphorylation of Ser and Thr in the NMBD does not change the interface of the NMII binding site, allosterically regulating the binding.

**Figure 8.**
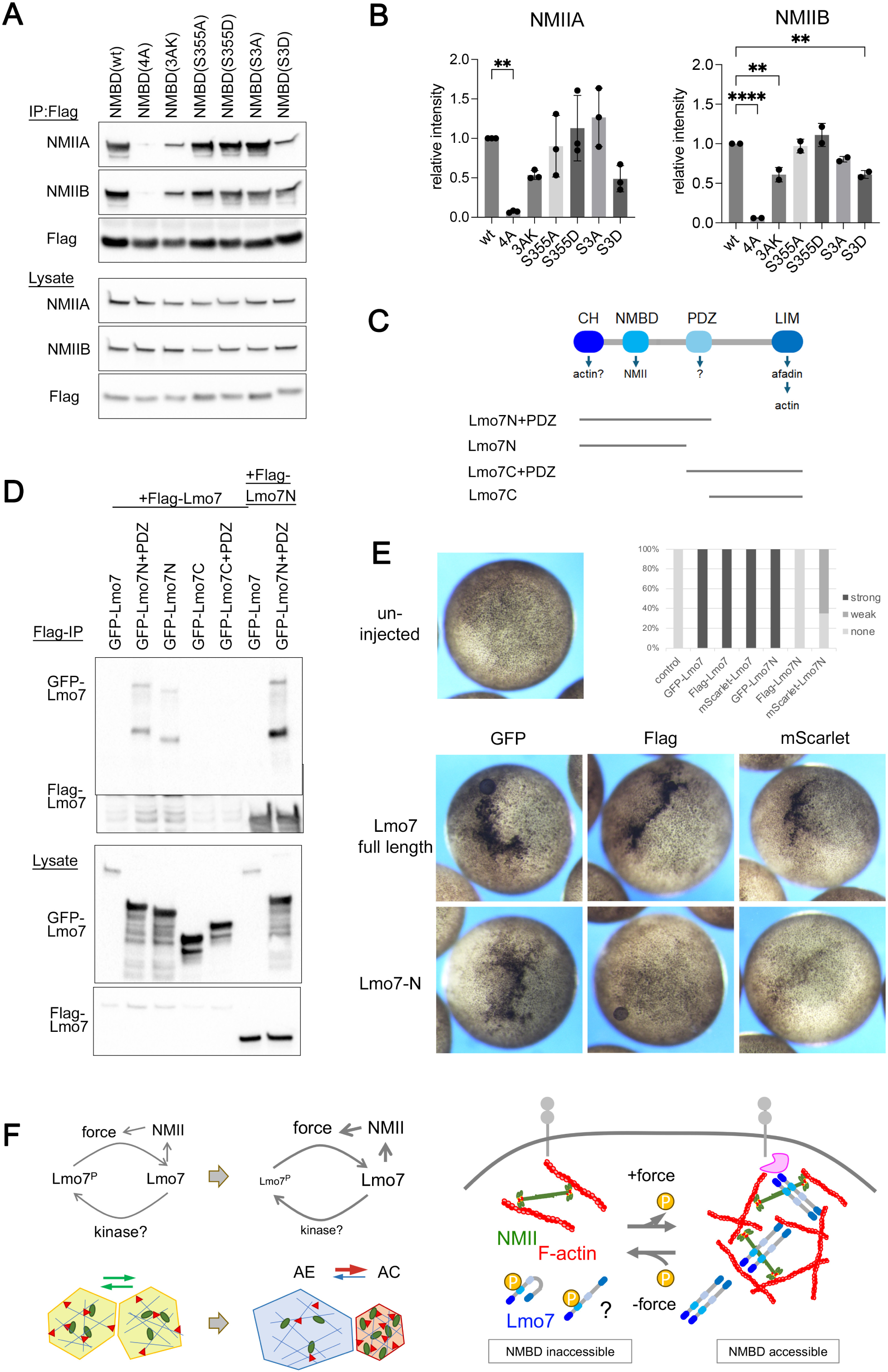
Lmo7 dimer formation and potential effects on the conformation changes for NMII binding. (A) Co-immunoprecipitation (co-IP) assays were conducted between NMIIA and the NMBD fragments of Lmo7 in HEK293T cells. The Flag-NMBD fragments included wild-type (wt) and several mutations: 4A, 3AK, S355D, S355A, S3D, and S3A. The co-IP products were analyzed using anti-NMIIA and anti-NMIIB antibodies. (B) The efficacy of NMIIA co-IP by Flag-NMBD was quantified from three independent experiments. For normalization, the band intensities from the anti-NMIIA IB were divided by those from the anti-Flag IB, which was further normalized to the intensity of Lmo7(wt). (C) Schematics of the Lmo7 deletion mutants used in the co-IP assays. (D) A co-IP assay was performed between Lmo7 deletion mutants in HEK293T cells. The co-IP products, obtained via anti-Flag IP were analyzed using an anti-GFP antibody. (E) Representative images of gastrula stage embryos overexpressing high levels of exogenous GFP-Lmo7, Flag-Lmo7, GFP-Lmo7N, and Flag-Lmo7N are shown. One nanogram of RNA was injected into diagonally located dorsal and ventral animal blastomeres in 4- or 8-cell stage embryos. The distribution of pigment granules in the animal side of the ectoderm was assessed at stage 11. The number of embryos analyzed includes: 32 for control, 22 for GFP-Lmo7, 37 for Flag-Lmo7, 22 for mScarlet-Lmo7, 74 for GFP-Lmo7N, 55 for Flag-Lmo7N, and 20 for mScarlet-Lmo7N. Scale bars: 50 μm. (F) A model for how feedback regulation between Lmo7 and actomyosin forces generates AC and AE cells through force-sensitive Ser-Thr phosphorylation in the NMBD, resulting in changes in Lmo7-NMII binding and Lmo7-mediated actomyosin network formation.

We next examined what could be the conformational change in Lmo7 by the phosphorylation of Ser and Thr in the NMBD. Lmo7 conformation may switch between closed and open forms. Alternatively, Lmo7 may form a dimer, altering the affinity to NMII. To test these possibilities, we conducted co-IP experiments using Lmo7 deletion mutants. While the interaction between full-length Lmo7 proteins was not confirmed, Flag-Lmo7 co-immunoprecipitated the N-terminal fragments of GFP-Lmo7, but not the C-terminal fragments (Fig. 8C, D). Flag-Lmo7N was sufficient to pull down GFP-Lmo7N (Fig. 8D). These results suggest that Lmo7 forms an oligomer through the N-terminal domain. Our previous study showed that GFP-Lmo7N promotes the formation of actomyosin network and induces collective AC (Matsuda et al., 2022). Notably, while Flag-Lmo7N binds NMII heavy chains in the co-IP assay (Matsuda et al., 2022), Flag-Lmo7N OE did not induce the accumulation of pigment granules or epithelial indentation (Fig. 8E), well-known indicators of coordinated AC in the *Xenopus* gastrula ectoderm (Haigo et al., 2003; Hildebrand, 2005). The mScarlet tag, which does not form oligomers (Bindels et al., 2017), also reduced the intensity of Lmo7-mediated coordinated AC, while the effect was less significant than that observed with the Flag tag (Fig. 8E, F). The activity of full-length Lmo7 in inducing coordinated AC was unaffected by either the Flag or mScarlet tags (top panels in Fig. 8E). These results suggest that the GFP tag at the N-terminus imparts additional changes to Lmo7N, either in its conformation or through post-translational modifications. This alteration appears to be critical for enhancing the activity of Lmo7 in increasing actomyosin contractility during AC..

## Discussion

This study shows that Lmo7 is a force-sensitive regulator of actomyosin in neuroectoderm during AC. High actomyosin contractility of neuroectoderm cells leads to junction shortening and the isotropic reduction of the apical domain. Whereas a subset of cells undergoes isotropic AC, the adjacent cells exhibit preferential elongation along the AP axis, resulting in the non-uniform reduction of their apical domain. This anisotropic AC is crucial for preserving the length of the NT during the folding process.

We showed that initially homogeneous Lmo7 overexpression in non-neural ectoderm progressively resulted in apical domain heterogeneity, suggesting that Lmo7 plays an important role in regulating anisotropic AC at the onset of NT folding. Pulsatile actomyosin activity at the apical cortex generates mutual pulling forces between adjacent cells, leading to fluctuations in actomyosin contractility and apical domain size. In our model, the positive feedback between Lmo7 and actomyosin force amplifies initially small differences in apical domain size between neighboring cells, leading to heterogeneity of apical domain size, shape and cell orientation. At the same time, the abundance of Lmo7 and actomyosin at the apical cortex is altered, so that the cells with smaller apical domain exhibit higher levels of Lmo7 at the apical surface (Left panels in Fig. 8F). We propose that this heterogeneity of the apical domain, facilitated by Lmo7, is essential for enabling cells in the neuroectoderm to elongate along the AP axis.

Mechanistically, phosphomimetic mutations at Ser355, Thr357, and Ser358 within the NMBD reduced NMII binding to full-length Lmo7, although this effect was less detectable in the NMBD fragment. These findings suggest that mechanical force leads to dephosphorylation of the conserved Ser and Thr in the NMBD, causing enhanced Lmo7 binding to NMII. Additionally, substituting alanine for the conserved WQ-WK sequence (a.a. 342, 343, 349, and 350) in the NMBD significantly suppressed NMII binding to both full-length Lmo7 and the NMBD fragment.

This, despite the close proximity of the WQ-WK sequence to the conserved Ser and Thr within the NMBD, S3D mutations probably interfere with the accessibility of NMII binding site due to a conformational change in the full-length Lmo7, while the 4A mutations directly alter the interface of the NMII binding site in the NMBD. The requirement of conformation changes for Lmo7 controlling actomyosin was further supported by the intermolecular interaction between Lmo7 in the co-IP assay and the requirement of GFP tag for the N-terminal fragment of Lmo7 to induce coordinated AC in *Xenopus* gastrula ectoderm. In our model, Lmo7 forms oligomer, which modulates Lmo7 association with NMII and F-actin, facilitating Lmo7 function on actomyosin (right panel in Fig. 8G). Additional studies and structural insights will be essential to explain the distinct effects of these mutations on the Lmo7 binding to NMII.

Different mechanisms have been described for the actomyosin control of AC in vertebrate and invertebrate epithelia. In invertebrates, adherens junctions lack well-developed actomyosin bundles. Instead, pulsatile actomyosin at the apical cortex generates the contractile forces necessary for AC through a “ratchet” mechanism (Martin and Goldstein, 2014; Martin et al., 2009; Miao and Blankenship, 2020). In contrast, vertebrate epithelia often feature circumferential actomyosin bundles beneath the apical junctions, which are thought to generate contractile forces in a “purse-string” manner. Recent studies, including ours, have shown that medial actomyosin pulsates dynamically in vertebrate epithelia, with increased abundance at the apical cortex during coordinated AC (Baldwin et al., 2023; Butler and Wallingford, 2018; Lardennois et al., 2019; Maitre et al., 2015; Shindo et al., 2019; Zhou et al., 2015). We observed that Lmo7 binding to NMII enhances the formation of both junctional and medioapical actomyosin networks. Therefore, it is likely that two apically localized actomyosin networks cooperatively regulate AC in vertebrate epithelia, but the role of Lmo7 in this regulation remains to be clarified in further studies.

While high levels of Lmo7 OE induced a sarcomere-like pattern in the actomyosin bundles underlying junctions (Matsuda et al., 2022), lower levels of Lmo7 OE, more relevant to the physiological Lmo7 function, resulted in a more significant increase in medial rather than junctional actomyosin during AC in the *Xenopus* gastrula ectoderm. In these AC cells, Lmo7 forms clusters at the apical cortex, immobilizing pulsatile actomyosin networks for extended periods, compared to the AC cells induced by Shroom3. Although the mechanism by which Lmo7 is recruited to the apical cortex remains unclear, we propose that Lmo7 acts as an adaptor, linking the actomyosin meshwork to the apical cortex. This connection would help stabilize AC by reducing or even terminating the oscillatory cycles of actomyosin contraction and relaxation between neighboring cells.

## Methods

### Plasmid, morpholino, and RNA preparation

pCS107-GFP-*Xenopus* Lmo7, pCS107-Flag-xLmo7, pCS107-Flag-xLmo7(4A), pCS2-GFP-NMIIA, pCS2-mScarlet-UtrCH, pCS2-lifeact-mScarlet, pCS2-lifeact-mNeon, pCS2-myr-tagBFP-HA, pCS2-myr-mRFP, pCS2-tagBFP were described in previous studies (Matsuda et al., 2022; Matsuda et al., 2023). Phospho-mimetic or non-phospho mutant forms of Lmo7 was generated using a standard *in vitro* mutagenesis protocol using KOD turbo DNA polymerase (TOYOBO) and the primers listed in supplemental information. pCS107-GFP-xPk2 was kindly provided by John Wallingford (University of Texas)(Butler and Wallingford, 2015; Butler and Wallingford, 2018). pCS2-mNeon-Sf9 were generous gifts from Ed Munro (University of Chicago, USA)(Hashimoto et al., 2015). pCS2-myc-*Xenopus* Shroom3 was described in a previous study (Ossipova et al., 2014). After linearizing plasmids by restriction enzymes, capped mRNA was synthesized using an mMessage mMachine SP6 Transcription Kit (ThermoFisher) and purified with an RNeasy Mini Kit (Qiagen). Lmo7 splicing blocking morpholino (MO) have been characterized in a previous study (GeneTools)(Matsuda et al., 2022)

### Cell culture, transfection, Immunoprecipitation and immunoblotting

HEK293T were purchased from ATCC and maintained in Dulbecco’s modified Eagle’s medium (DMEM)(Corning) supplemented with 10% fetal bovine serum (Sigma) in a 37 C incubator with 5% CO2. HEK293T cells grown in 60 mm dish to 70-80% density were transfected with a mixture of 4 μg plasmid DNA and 20 μg polyethylenimine (Polysciences). Cell lysates were extracted 24 hrs after transfection using the Radioimmunoprecipiration (RIPA) buffer (50 mM Tris-HCl pH 7.4, 150 mM NaCl, 1% NP-40, 0.5% sodium deoxycholate, 0.1% SDS), supplemented with Protease Inhibitor Cocktail III (Calbiochem) and PhosStop phosphatase inhibitor cocktail (Roche). Cell lysates were incubated with anti-Flag M2 agarose (Sigma) at 4°C at least 6 hours. Agarose beads were washed three times in Tris-buffered saline containing 0.05% Triton X-100 (TBST). Agarose beads were heated at 95°C for 5 min in SDS sample buffer containing 5% β-mercaptoethanol (Sigma). After SDS-PAGE and transfer to a nitrocellulose membrane with a 0.2 μm pore size (RioRad), western blots were performed. Antibodies used are the following: rabbit anti-NMIIA pAb (BioLegend, #POLY19098, 1:500) and rabbit anti-NMIIB pAb (BioLegend, #POLY19099, 1:500), mouse anti-DYKDDDDK mAb clone 2H8 (Cosmo Bio USA, #KAL-K0602, 1:1000). HRP-conjugated secondary antibodies against mouse or rabbit IgG were from Cell Signaling Technology (#7074, #7076, 1:5000). Chemiluminescent signals were acquired using Clarity ECL Western Blotting Substrates (Bio-Rad) on the ChemiDoc MP Imaging System (Bio-Rad).

### Xenopus embryos and microinjection

Wild-type *Xenopus laevis* were purchased from Nasco and Xenopus1, maintained and handled following the Guide for the Care and Use of Laboratory Animals of the National Institutes of Health. The protocol for animal use was approved by the Institutional Animal Care and Use Committee (IACUC) of the Icahn School of Medicine at Mount Sinai. The sex of the animals was not considered in the study design or analysis because the study subjects were sexually indifferent embryos. *In vitro* fertilization and embryo culture were performed as previously described (Ossipova et al., 2014). Embryo staging was determined according to Nieuwkoop and Faber (Nieuwkoop and Faber, 1994). For microinjections, embryos were transferred to 3% Ficoll 400 (Pharmacia) in 0.5× Marc’s modified Ringer’s (MMR) solution (50 mM NaCl, 1 mM KCl, 1 mM CaCl2, 0.5 mM MgCl2 and 2.5 mM HEPES (pH 7.4))(Peng, 1991). *in vitro* synthesized RNA or MO in 5-10 nl of RNase-free water (Invitrogen) was microinjected into one to two animal blastomeres of 4-8 cell-stage embryos. For the imaging of apical domain dynamics in the ectoderm of the gastrula-stage embryos, two ventral-animal blastomeres were injected with 150 pg of RNA encoding GFP-Lmo7, coinjected with 50 pg of RNA encoding myr-tagBFP-HA or myr-mRFP. For the imaging of actomyosin dynamics in the gastrula ectoderm, 300 pg of Flag-Lmo7 RNA or 100 pg of myc-Shroom3 RNA was injected to two ventral-animal blastomeres, with 100 pg of sf9-mNeon and 100 pg of lifeact-mScarlet or 50 pg of Scarlet-UtrCH RNA. For the imaging of apical domain dynamics in the neuroectoderm, 20 ng of Lmo7 MO was injected into one dorsal-animal blastomere with 50 pg of lifeact-mNeon RNA. 100 pg of lifeact-Scarlet RNA was injected into the other dorsal-animal blastomere in the same embryos. For the imaging of actomyosin dynamics in the neuroectoderm, 100 pg of mNeon-Sf9 RNA and 100 pg of lifeact-Scarlet RNA were coinjected in two dorsal-animal blastomeres, followed by the injection of 20 ng of Lmo7 MO with tracer tagBFP RNA in one dorsal-animal blastomere. For imaging to determine Pk2 distribution, 100 pg of GFP-Pk2 RNA was coinjected with 100 pg of lifeact-mScarlet RNA into one dorsal-animal blastomere, with and without 20 ng of Lmo7 MO or 10 ng of Vangl2 MO. RNA- or MO-injected embryos were cultured in 0.1x MMR until the early gastrula or neurula stage. Each injection included at least 20 embryos per condition. The experiments were repeated at least three times.

### phalloidin staining, immunofluorescence, and imaging of fixed Xenopus embryos

For phalloidin staining, devitellinized embryos were fixed in MEMFA (100 mM MOPS (pH 7.4), 2 mM EGTA, 1 mM MgSO_4_, 3.7% formaldehyde)(Harland, 1991) for 1 hr at room temperature. After permeabilization in 0.1% Triton X-100 in PBS for 10 min, the embryos were incubated with Alexa Fluor 555-conjugated phalloidin (1:400 dilution, ThermoFisher Scientific) in PBS containing 1% BSA overnight at 4°C. For immunostaining, devitellinized embryos were fixed in ice-cold 10% Trichloroacetic acid (TCA) for 30 min. After washing several times in TBS, embryos were incubated in 1% BSA in TBS at 4°C overnight for blocking. The incubation of primary and secondary antibodies was in 1% BSA and 1% DMSO in TBS at 4°C overnight. Antibodies used in this study are the following: rabbit anti-NMIIA pAb (BioLegend, POLY19098, 1:300), rat anti-HA mAb clone 3F10 (Roche, 118673230002, 1:100), and chicken anti-GFP pAb (abcam, ab13970, 1:500), Alexa 488-conjugated anti-chicken IgY (Jackson ImmunoResearch) and Cy3-conjugated anti-rabbit IgG (Jackson ImmunoResearch Laboratory #711-165-152, 1:500). AlexaFluor 555-phalloidin (Invitrogen, A34055) was added to secondary antibody solution to label F-actin. After wash in PBS or TBS three times, embryos were transferred in 25% glycerol in PBS. The dissected neural plate was mounted on a glass slide with two coverglass spacers (0.13–0.17 mm) to minimize damage to the morphology of the neuroectoderm.

Images of fixed and stained embryos were captured either by a BC43 spinning disk confocal microscope (Fusion Ver 2, Andor, Oxford Instruments) or a Zeiss LSM980 with an Airyscan 2 confocal microscope (Zen (blue edition) Ver 3.7.97.04000, Zeiss). The Nikon CFI Plan Apochromat Lambda D 20X (NA=0.8 and WD=0.8 mm) or Zeiss Plan-Apochromat 20x/0.8 (NA=0.8 and WD=0.55 mm) were used for image acquisition. The tile function of Fusion and Zen was used to stitch images. Z-stack images were projected into a single image via the maximum projection function in ImageJ2 (ver. 2.9.0/1.54g), which was used for further analysis and quantification. Each experiment included at least 3 embryos per experimental group, and the experiments were repeated at least three times.

### Time-lapse imaging of live Xenopus embryos

Five to ten embryos per individual group were mounted in 1% low melting temperature agarose (SeaPlaque agarose, #50101, Lonza) in 0.1× MMR on a glass slide attached to a silicone isolator (1.2 mm, Grace Biolabs) or on a glass-bottom dish (#1, Cellvis). Time-lapse imaging was carried out at room temperature either on an Andor BC43 spinning disk confocal microscope, or a Zeiss AxioZoomV16 fluorescence stereomicroscope equipped with an AxioCam 506 mono CCD camera. A Nikon CFI Plan Apochromat Lambda D 20X (NA=0.8 and WD=0.8 mm) was used for the imaging to assess the apical domain dynamics. Nikon CFI Plan Apo Lambda S 40XC Sil (NA=1.25 and WD=0.3 mm) was used for the imaging to assess actomyosin dynamics. Images were taken every 1 second to 5 minutes for a period of 15 min to 5 hrs. The multiposition tool in Fusion or Zen (blue) imaging software was used for simultaneous time-lapse imaging. Z-stack images were projected into a single image via the maximum projection function in ImageJ. Each experiment included at least 3 embryos per experimental group, and the experiments were repeated at least three times.

### Data analysis, segmentation, cell tracking, and the assessment of apical domains and junctions

Image processing and quantification were performed as previously described (Matsuda et al., 2023; Matsuda and Sokol, 2025). Briefly, grayscale images of the cell outline marker were segmented via the Python package CellPose v2.0.5 (Pachitariu and Stringer, 2022). Segmented cells were tracked across timepoints via the Python package Bayesian Tracker (btrack v0.4.5)(Ulicna et al., 2021) with manual correction. The border of the neuroectoderm was defined by the regions with brighter signals of Lifeact-mNeon, Lifeact-mScarlet or phalloidin than those in the surrounding non-neural ectoderm. After segmentation, each tricellular junction (TCJ) was first identified as a vertex or edge of the cell outline network. Bicellular junction (BJ) was identified between two vertices. The mean fluorescence intensities were measured via dilated masks of the pixel or a set of pixels that defined BJs and TCJs (0.76155 μm in width or diameter). The medioapical domain was the area that did not correspond to TCJs or BJs.

### Statistics & Reproducibility

Histograms and dot plots of the experimental data were generated via GraphPad Prism 10 (ver. 10.1.0) and Microsoft Excel. Individual experiments were repeated at least three times. The standard deviations (S.D.) were calculated via GraphPad Prism 10 and used to assess the variability among sample groups. Student’s t test or the Mann‒Whitney test was used to compare means or ranks between two groups, respectively. One-way ANOVA was used to compare the means of more than two groups, with the Bonferroni correction for pairwise comparisons. The two-sample Kolmogorov‒Smirnov test was used to compare distributions between two groups. The chi-square test was used for the categorical data.

## Acknowledgments

We thank L. Davidson, A. Miller, E. Munro, and J. Wallingford for plasmids. We acknowledge the help from the ISMMS Microscopy Core facility. This research was supported by the NIH grant R35GM122492 to S.Y.S.

## Author contribution statement

M.M. and S.Y.S. conceptualized and developed the project. M.M. designed and performed the experiments and the formal analyses. M.M. and S.Y.S. contributed to the data interpretation. M.M. prepared the figures and wrote the original manuscript. M.M. and S.Y.S. edited the manuscript. Both authors approved the final manuscript.

## References

Baldwin, A., I.K. Popov, R. Keller, J. Wallingford, and C. Chang. 2023. The RhoGEF protein Plekhg5 regulates medioapical and junctional actomyosin dynamics of apical constriction during Xenopus gastrulation. Mol Biol Cell. 34:ar64.

Baldwin, A.T., J.H. Kim, H. Seo, and J.B. Wallingford. 2022. Global analysis of cell behavior and protein dynamics reveals region-specific roles for Shroom3 and N-cadherin during neural tube closure. Elife. 11.

Bardet, P.L., B. Guirao, C. Paoletti, F. Serman, V. Leopold, F. Bosveld, Y. Goya, V. Mirouse, F. Graner, and Y. Bellaiche. 2013. PTEN controls junction lengthening and stability during cell rearrangement in epithelial tissue. Dev Cell. 25:534–546.

Baum, B., J. Settleman, and M.P. Quinlan. 2008. Transitions between epithelial and mesenchymal states in development and disease. Semin Cell Dev Biol. 19:294–308.

Beati, H., I. Peek, P. Hordowska, M. Honemann-Capito, J. Glashauser, F.A. Renschler, P. Kakanj, A. Ramrath, M. Leptin, S. Luschnig, S. Wiesner, and A. Wodarz. 2018. The adherens junction-associated LIM domain protein Smallish regulates epithelial morphogenesis. J Cell Biol. 217:1079–1095.

Benton, M.A., N. Frey, R. Nunes da Fonseca, C. von Levetzow, D. Stappert, M.S. Hakeemi, K.H. Conrads, M. Pechmann, K.A. Panfilio, J.A. Lynch, and S. Roth. 2019. Fog signaling has diverse roles in epithelial morphogenesis in insects. Elife. 8.

Bindels, D.S., L. Haarbosch, L. van Weeren, M. Postma, K.E. Wiese, M. Mastop, S. Aumonier, G. Gotthard, A. Royant, M.A. Hink, and T.W. Gadella, Jr. 2017. mScarlet: a bright monomeric red fluorescent protein for cellular imaging. Nat Methods. 14:53–56.

Butler, M.T., and J.B. Wallingford. 2015. Control of vertebrate core planar cell polarity protein localization and dynamics by Prickle 2. Development. 142:3429–3439.

Butler, M.T., and J.B. Wallingford. 2018. Spatial and temporal analysis of PCP protein dynamics during neural tube closure. Elife. 7.

Christodoulou, N., and P.A. Skourides. 2015. Cell-Autonomous Ca(2+) Flashes Elicit Pulsed Contractions of an Apical Actin Network to Drive Apical Constriction during Neural Tube Closure. Cell Rep. 13:2189–2202.

Christodoulou, N., and P.A. Skourides. 2022. Distinct spatiotemporal contribution of morphogenetic events and mechanical tissue coupling during Xenopus neural tube closure. Development. 149.

Ciruna, B., A. Jenny, D. Lee, M. Mlodzik, and A.F. Schier. 2006. Planar cell polarity signalling couples cell division and morphogenesis during neurulation. Nature. 439:220–224.

Colas, J.F., and G.C. Schoenwolf. 2001. Towards a cellular and molecular understanding of neurulation. Dev Dyn. 221:117–145.

Copp, A.J., and N.D. Greene. 2010. Genetics and development of neural tube defects. J Pathol. 220:217–230.

Costa, M., E.T. Wilson, and E. Wieschaus. 1994. A putative cell signal encoded by the folded gastrulation gene coordinates cell shape changes during Drosophila gastrulation. Cell. 76:1075–1089.

Das, D., J.K. Zalewski, S. Mohan, T.F. Plageman, A.P. VanDemark, and J.D. Hildebrand. 2014. The interaction between Shroom3 and Rho-kinase is required for neural tube morphogenesis in mice. Biol Open. 3:850–860.

Du, T.T., J.B. Dewey, E.L. Wagner, R. Cui, J. Heo, J.J. Park, S.P. Francis, E. Perez-Reyes, S.J. Guillot, N.E. Sherman, W. Xu, J.S. Oghalai, B. Kachar, and J.B. Shin. 2019. LMO7 deficiency reveals the significance of the cuticular plate for hearing function. Nat Commun. 10:1117.

Eiraku, M., N. Takata, H. Ishibashi, M. Kawada, E. Sakakura, S. Okuda, K. Sekiguchi, T. Adachi, and Y. Sasai. 2011. Self-organizing optic-cup morphogenesis in three-dimensional culture. Nature. 472:51–56.

Francou, A., K.V. Anderson, and A.K. Hadjantonakis. 2023. A ratchet-like apical constriction drives cell ingression during the mouse gastrulation EMT. Elife. 12.

Haigo, S.L., J.D. Hildebrand, R.M. Harland, and J.B. Wallingford. 2003. Shroom induces apical constriction and is required for hingepoint formation during neural tube closure. Curr Biol. 13:2125–2137.

Harland, R.M. 1991. In situ hybridization: an improved whole-mount method for Xenopus embryos. Methods Cell Biol. 36:685–695.

Hashimoto, H., F.B. Robin, K.M. Sherrard, and E.M. Munro. 2015. Sequential contraction and exchange of apical junctions drives zippering and neural tube closure in a simple chordate. Dev Cell. 32:241–255.

Hashimoto, Y., N. Kinoshita, T.M. Greco, J.D. Federspiel, P.M. Jean Beltran, N. Ueno, and I.M. Cristea. 2019. Mechanical Force Induces Phosphorylation-Mediated Signaling that Underlies Tissue Response and Robustness in Xenopus Embryos. Cell Syst. 8:226–241 e227.

Hildebrand, J.D. 2005. Shroom regulates epithelial cell shape via the apical positioning of an actomyosin network. J Cell Sci. 118:5191–5203.

Hildebrand, J.D., and P. Soriano. 1999. Shroom, a PDZ domain-containing actin-binding protein, is required for neural tube morphogenesis in mice. Cell. 99:485–497.

Humphries, A.C., and M. Mlodzik. 2018. From instruction to output: Wnt/PCP signaling in development and cancer. Curr Opin Cell Biol. 51:110–116.

Ikawa, K., S. Ishihara, Y. Tamori, and K. Sugimura. 2023. Attachment and detachment of cortical myosin regulates cell junction exchange during cell rearrangement in the Drosophila wing epithelium. Curr Biol. 33:263–275 e264.

Lardennois, A., G. Pasti, T. Ferraro, F. Llense, P. Mahou, J. Pontabry, D. Rodriguez, S. Kim, S. Ono, E. Beaurepaire, C. Gally, and M. Labouesse. 2019. An actin-based viscoplastic lock ensures progressive body-axis elongation. Nature. 573:266–270.

Maitre, J.L., R. Niwayama, H. Turlier, F. Nedelec, and T. Hiiragi. 2015. Pulsatile cell-autonomous contractility drives compaction in the mouse embryo. Nat Cell Biol. 17:849–855.

Manning, A.J., K.A. Peters, M. Peifer, and S.L. Rogers. 2013. Regulation of epithelial morphogenesis by the G protein-coupled receptor mist and its ligand fog. Sci Signal. 6:ra98.

Martin, A.C., and B. Goldstein. 2014. Apical constriction: themes and variations on a cellular mechanism driving morphogenesis. Development. 141:1987–1998.

Martin, A.C., M. Kaschube, and E.F. Wieschaus. 2009. Pulsed contractions of an actin-myosin network drive apical constriction. Nature. 457:495–499.

Matsuda, M., C.W. Chu, and S.Y. Sokol. 2022. Lmo7 recruits myosin II heavy chain to regulate actomyosin contractility and apical domain size in Xenopus ectoderm. Development. 149.

Matsuda, M., J. Rozman, S. Ostvar, K.E. Kasza, and S.Y. Sokol. 2023. Mechanical control of neural plate folding by apical domain alteration. Nat Commun. 14:8475.

Matsuda, M., and S.Y. Sokol. 2025. Prickle2 regulates apical junction remodeling and tissue fluidity during vertebrate neurulation. J Cell Biol. 224.

Miao, H., and J.T. Blankenship. 2020. The pulse of morphogenesis: actomyosin dynamics and regulation in epithelia. Development. 147.

Nieuwkoop, P.D., and J. Faber. 1994. Normal table of Xenopus laevis (Daudin): a systematical and chronological survey of the development from the fertilized egg till the end of metamorphosis. Garland Pub., New York. 252 p., 210 leaves of plates pp.

Ooshio, T., K. Irie, K. Morimoto, A. Fukuhara, T. Imai, and Y. Takai. 2004. Involvement of LMO7 in the association of two cell-cell adhesion molecules, nectin and E-cadherin, through afadin and alpha-actinin in epithelial cells. J Biol Chem. 279:31365–31373.

Ossipova, O., K. Kim, B.B. Lake, K. Itoh, A. Ioannou, and S.Y. Sokol. 2014. Role of Rab11 in planar cell polarity and apical constriction during vertebrate neural tube closure. Nat Commun. 5:3734.

Pachitariu, M., and C. Stringer. 2022. Cellpose 2.0: how to train your own model. Nat Methods.

Peng, H.B. 1991. Xenopus laevis: Practical uses in cell and molecular biology. Solutions and protocols. Methods Cell Biol. 36:657–662.

Peng, Y., and J.D. Axelrod. 2012. Asymmetric protein localization in planar cell polarity: mechanisms, puzzles, and challenges. Curr Top Dev Biol. 101:33–53.

Perez-Vale, K.Z., and M. Peifer. 2020. Orchestrating morphogenesis: building the body plan by cell shape changes and movements. Development. 147.

Ramkumar, N., T. Omelchenko, N.F. Silva-Gagliardi, C.J. McGlade, J. Wijnholds, and K.V. Anderson. 2016. Crumbs2 promotes cell ingression during the epithelial-to-mesenchymal transition at gastrulation. Nat Cell Biol. 18:1281–1291.

Sawyer, J.M., J.R. Harrell, G. Shemer, J. Sullivan-Brown, M. Roh-Johnson, and B. Goldstein. 2010. Apical constriction: a cell shape change that can drive morphogenesis. Dev Biol. 341:5–19.

Schoenwolf, G.C., and J.L. Smith. 1990. Mechanisms of neurulation: traditional viewpoint and recent advances. Development. 109:243–270.

Sherrard, K., F. Robin, P. Lemaire, and E. Munro. 2010. Sequential activation of apical and basolateral contractility drives ascidian endoderm invagination. Curr Biol. 20:1499–1510.

Shindo, A., Y. Inoue, M. Kinoshita, and J.B. Wallingford. 2019. PCP-dependent transcellular regulation of actomyosin oscillation facilitates convergent extension of vertebrate tissue. Dev Biol. 446:159–167.

Suzuki, M., M. Sato, H. Koyama, Y. Hara, K. Hayashi, N. Yasue, H. Imamura, T. Fujimori, T. Nagai, R.E. Campbell, and N. Ueno. 2017. Distinct intracellular Ca(2+) dynamics regulate apical constriction and differentially contribute to neural tube closure. Development. 144:1307–1316.

Uechi, H., and E. Kuranaga. 2019. The Tricellular Junction Protein Sidekick Regulates Vertex Dynamics to Promote Bicellular Junction Extension. Dev Cell. 50:327–338 e325.

Ulicna, K., G. Vallardi, G. Charras, and A.R. Lowe. 2021. Automated Deep Lineage Tree Analysis Using a Bayesian Single Cell Tracking Approach. Frontiers in Computer Science. 3.

Vielemeyer, O., C. Nizak, A.J. Jimenez, A. Echard, B. Goud, J. Camonis, J.C. Rain, and F. Perez. 2010. Characterization of single chain antibody targets through yeast two hybrid. BMC Biotechnol. 10:59.

Wallingford, J.B., L.A. Niswander, G.M. Shaw, and R.H. Finnell. 2013. The continuing challenge of understanding, preventing, and treating neural tube defects. Science. 339:1222002.

West, J.J., T. Zulueta-Coarasa, J.A. Maier, D.M. Lee, A.E.E. Bruce, R. Fernandez-Gonzalez, and T.J.C. Harris. 2017. An Actomyosin-Arf-GEF Negative Feedback Loop for Tissue Elongation under Stress. Curr Biol. 27:2260–2270 e2265.

Williams, M., C. Burdsal, A. Periasamy, M. Lewandoski, and A. Sutherland. 2012. Mouse primitive streak forms in situ by initiation of epithelial to mesenchymal transition without migration of a cell population. Dev Dyn. 241:270–283.

Ybot-Gonzalez, P., D. Savery, D. Gerrelli, M. Signore, C.E. Mitchell, C.H. Faux, N.D. Greene, and A.J. Copp. 2007. Convergent extension, planar-cell-polarity signalling and initiation of mouse neural tube closure. Development. 134:789–799.

Zhou, J., S. Pal, S. Maiti, and L.A. Davidson. 2015. Force production and mechanical accommodation during convergent extension. Development. 142:692–701.

